# OptoCRISPRi-HD: engineering a green-light activated CRISPRi system with high dynamic range

**DOI:** 10.1101/2022.12.31.522379

**Authors:** Ke-Ning Chen, Bin-Guang Ma

**Author notes:** Corresponding author. *E-mail address* (Bin-Guang Ma).

## Abstract

The ability to modulate gene expression is crucial for studying gene function and programming cell behaviors. Combining the reliability of CRISPRi and the precision of optogenetics, the opto-CRISPRi technique is emerging as an advanced tool for live-cell gene regulation. Since previous versions of opto-CRISPRi often exhibit a no more than 10 folds’ dynamic range due to the leakage activity, they are not suitable for targets that are sensitive to such leakage or critical for cell growth. Here, we describe a green-light activated CRISPRi system with high dynamic range (40-fold) and the flexibility of changing targets in *Escherichia coli*. Our optoCRISPRi-HD system can efficiently repress essential genes, non-essential genes or inhibit the initiation of DNA replication. Providing a regulative system with high resolution over space-time and extensive targets, our study would facilitate further researches involving complex gene networks, metabolic flux redirection or bioprinting.

## INTRODUCTION

Among all the inducers that are used in synthetic biology, light has been seen as the most advanced and promising one. Light owns some obvious advantages: it can be projected reversibly with spatial and temporal precision, divided into different wave lengths and programmed by a computer. It is logical and practical to choose light as the switch of CRISPRi (CRISPR interference) system^1^, since dCas9 shows cell toxicity^2^ and hence cannot be expressed constitutively. The combination of optogenetics and CRISPRi makes an ideal tool for multiple gene regulation and automatic cell behavior control.

Seeking to achieve a light controlled CRISPRi, several approaches have been made and tested by researchers. A blue-light induced CRISPRi was constructed by putting the expression of sgRNA under the control of YF1-FixJ system.^3^ Another blue-light controlled CRISPRi was made by fusing split dCas9 fragments with photoinducible dimerization domains.^4^ In a study of RGB light sensing system, CRISPRi was induced by red, green or blue light that controls sgRNA expression through three different two-component signal transduction systems (TCSs).^5^ In addition, a dCpf1 (also known as dCas12a) based blue-light activated CRISPRi system used EL222 as light sensor and transcriptional activator.^6^ These studies tend to use blue light as inducers, whereas it is widely understood that blue light can cause cell damage in both prokaryotes and eukaryotes.^7–9^ Meanwhile, these previous versions of opto-CRISPRi often exhibit a no more than 10 folds’ dynamic range due to the leakage activity before light induction. For targets that are sensitive to this leakage or critical for cell growth, the opto-CRISPRi may face the failure of “switching off” or lead to a severely influenced cell growth. The leakage activity of dCas9 can also impose a selective pressure upon the cell population to retain mutated plasmids instead of the functional ones.

In consideration of the issues above, we chose an optimized version of CcaS-CcaR TCS to control the expression of dCas9 and sgRNA. After being cloned into *E. coli* from *Synechocystis* PCC6803,^10^ CcaS-CcaR system went through some major improvements: The 238bp *P_cpcG2_* promoter was truncated into 172bp;^11^ The expression levels of CcaS, CcaR and Ho1-PcyA^12^ are optimized;^11^ Two PAS domains were deleted from CcaS.^13,14^ These improvements significantly reduced the leakage of CcaS-CcaR system and made it the best engineered green light sensor available. Specifically, green light (535nm) induces the CcaS phosphorylation and the phosphotransfer to CcaR, whereas red light (672nm) reverses the phosphorylation of CcaS and stops CcaR from receiving phosphate.^15^ In our design, phosphorylated and dimerized CcaR binds the *P_cpcG2_* promoter to activate the transcription of dCas9 and sgRNA, which leads to the silence of target gene under green light and a normal state under red light (**Figure 1**). By coupling CcaS-CcaR and CRISPRi, we aimed to develop a genome-wide optogenetic regulative system which is less harmful to bacteria than blue-light-induced CRISPRi and tight enough to control essential genes.

**Figure 1.**
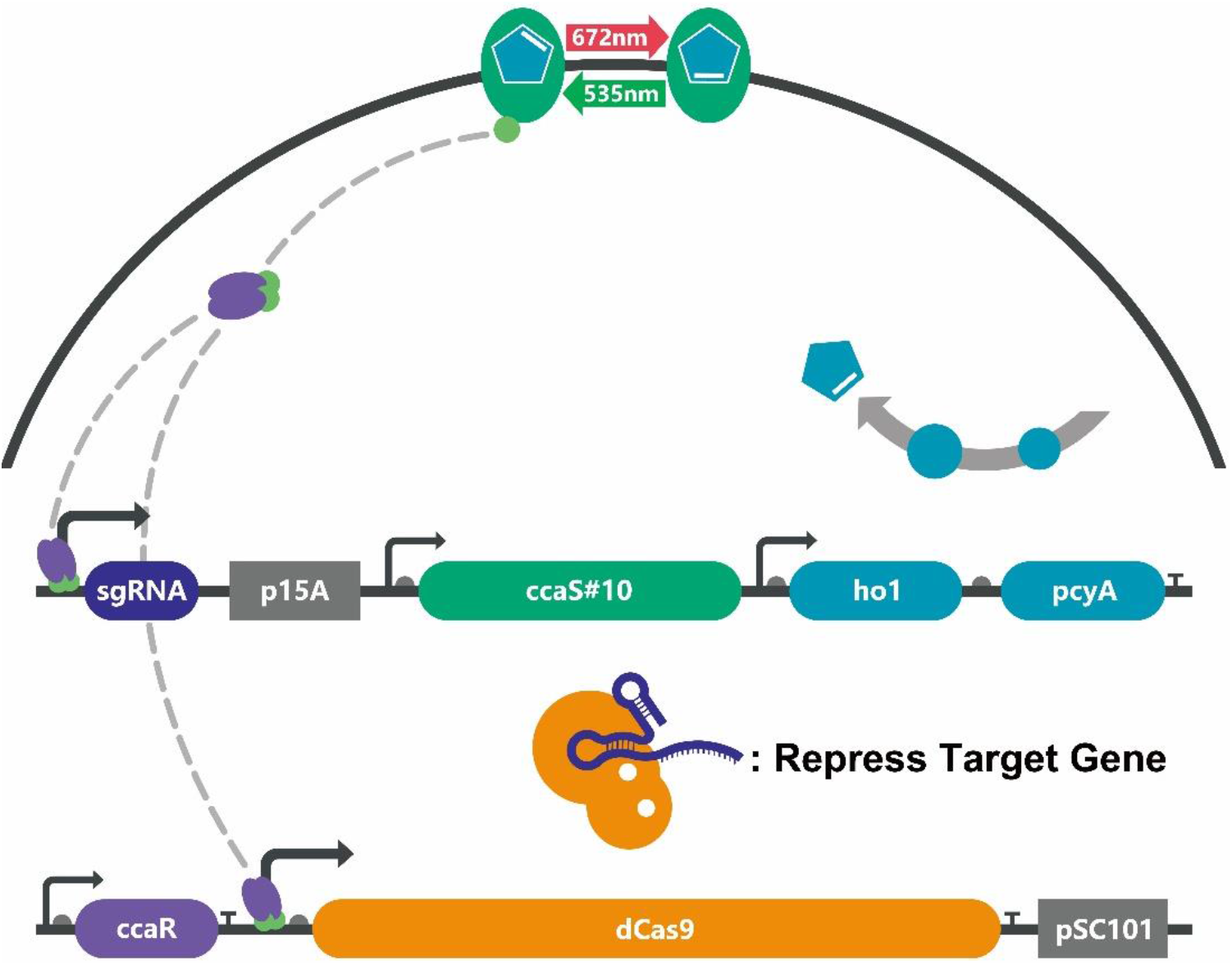
The circuit design and working mechanism of optoCRISPRi-HD.

## RESULTS

### Reducing the Leakage Expression of Opto-CRISPRi

In order to minimize the leakage expression under red light, we relocated the *ccaR* gene and *P_cpcG2_* promoter from a *ColE1-origin* plasmid (~50-70 copy)^11^ to a *pSC101-origin* plasmid (~1-5 copy) to lead dCas9 expression. The expression level of CcaR needed to be readjusted after this change. Using eGFP as reporter, we first replaced the original *ccaR* promoter *J23100* with four Anderson collection promoters of different strengths, among which the *J23117* showed the best performance of an 8.3-fold change between green/red light (**Figure 2A**). While keeping the *J23117* as promoter, we next tested a series of Ribosome Binding Sites (RBSs) to find the best dynamic range. Among these tested 7 RBSs, *J61110* showed the highest 32.3-fold change but the lowest expression level under green light (**Figure 2B**). In order to ensure the amount of dCas9 for a full repression, we chose the second highest 23.8-fold *J61108* as *ccaR* RBS for all the following experiments.

**Figure 2.**
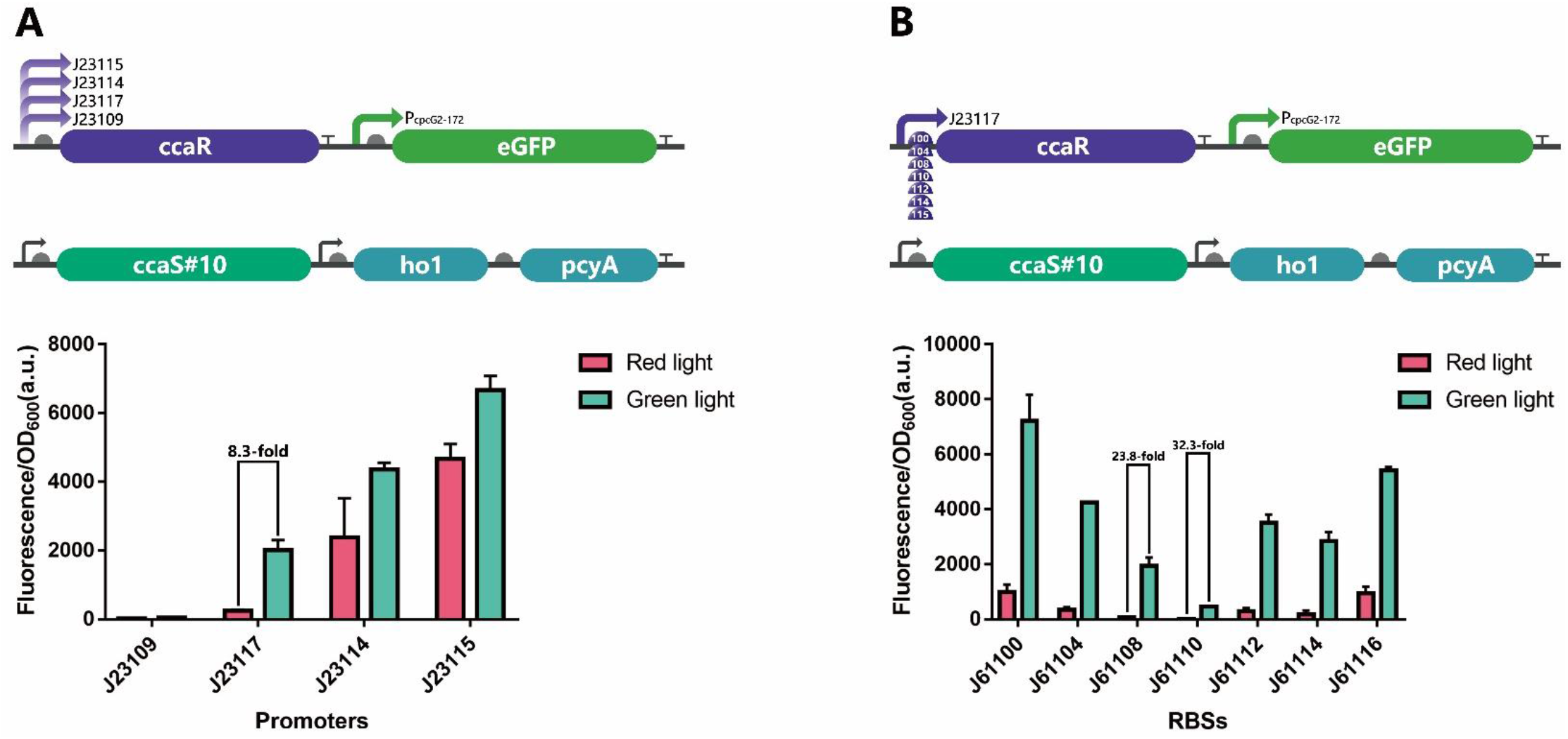
Reducing the leakage expression of *PcpcG2-172* by adjusting CcaR level. (A) Promoter changing experiments, among which *J23117* showed the highest 8.3-fold change between green/red light. (B) RBS changing experiments, among which *J61110* showed the highest 32.3-fold and J61108 the second highest 23.8-fold change between green/red light. All experiments were carried out with triplicate biological repeats.

After determining the expression level of CcaR, we put dCas9 and sgRNA separately on different plasmids, both controlled by *P_cpcG2-172_*. Two PAS domains were deleted from CcaS to obtain CcaS#10, which has been reported for having positive effect on green/red dynamic range.^14^ The final version of this system is named optoCRISPRi-HD (HD for high dynamic).

### Repressing *tetR-egfp* on *E. coli* Genome with optoCRISPRi-HD

For testing and quantifying the capacity of optoCRISPRi-HD, we constructed MGFROS strain based on *E. coli* K-12 MG1655. Fluorescent Repressor-Operator System (FROS) is developed to label DNA in live cell and observe the spatial position of interested loci in real time.^16^ As for our strain, a 120-repeat *tetO* sequence carrying gentamicin resistance gene was inserted into MG1655 genome near the origin of chromosome DNA replication (*oriC)*.The *tetR-eGFP* was inserted into genomic *ara* operon for a tight regulation (**Figure 3A**).

**Figure 3.**
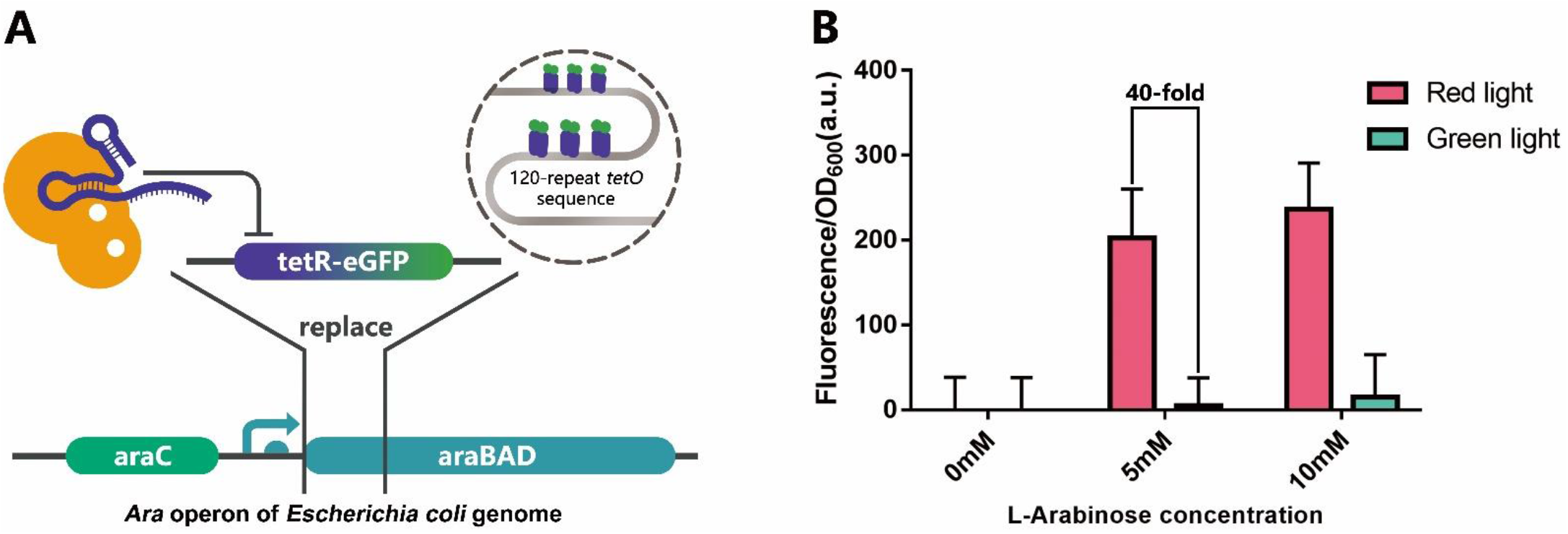
Using optoCRISPRi-HD to repress *tetR-eGFP* that was inserted into *E. coli* genome. (A) An illustration of the function and position of *tetR-eGFP*. (B) OptoCRISPRi-HD that targets *tetR-eGFP* created a 40-fold change between red/green in 5mM arabinose condition. The experiment was carried out with triplicate biological repeats.

Plasmid pvCq1 (carrying *tetR* sgRNA) was co-transformed with pvS108 (**Figure S1**) into MGFROS strain. The P_BAD_ controlled TetR-eGFP will begin to generate when arabinose is added into culture media. Under green light, the optoCRISPRi-HD is expressed to reduce TetR-eGFP, which leads to a much lower fluorescence value than red light condition. As shown in **Figure 3B**, our optoCRISPRi-HD achieved a 40-fold dynamic range against 5mM arabinose, which is much higher than most light controlled CRISPRi systems.

### Light Controlled Bioprinting by Repressing *lacZ*

After quantifying the dynamic range of optoCRISPRi-HD, we co-transformed plasmid pvCq7 (carrying *lacZ* sgRNA) and pvS108 (**Figure S1**) into *E. coli* Nissle1917 strain to perform a light controlled bioprinting. On an *E. coli* culture plate, β-galactosidase cleaves its substrate X-gal to appear a blue color. In this case, genomic β-galactosidase gene *lacZ* is repressed by optoCRISPRi-HD under green light and (almost) not influenced under red light, therefore leaving an image on the plate after green/red light exposure.

As shown in **Figure 4A**, two different strategies were applied to produce image: using a hollowed-out tin foil or a movie projector. A colony harboring pvCq7 and pvS108 was transferred into 1ml liquid LB and cultivated for 5 hours, 300μl of which was then subjected onto an IPTG-X-gal plate and put under light control. After 12 hours’ exposure, both strategies left traces on culture plates where the green light induced regions remained uncolored due to a lack of β-galactosidase (**Figure 4B**). This result indicated the robustness of optoCRISPRi-HD for the tolerance of inaccurate wavelengths (In this case, the movie projector can only produce an approximate wavelength of green or red light).

**Figure 4.**
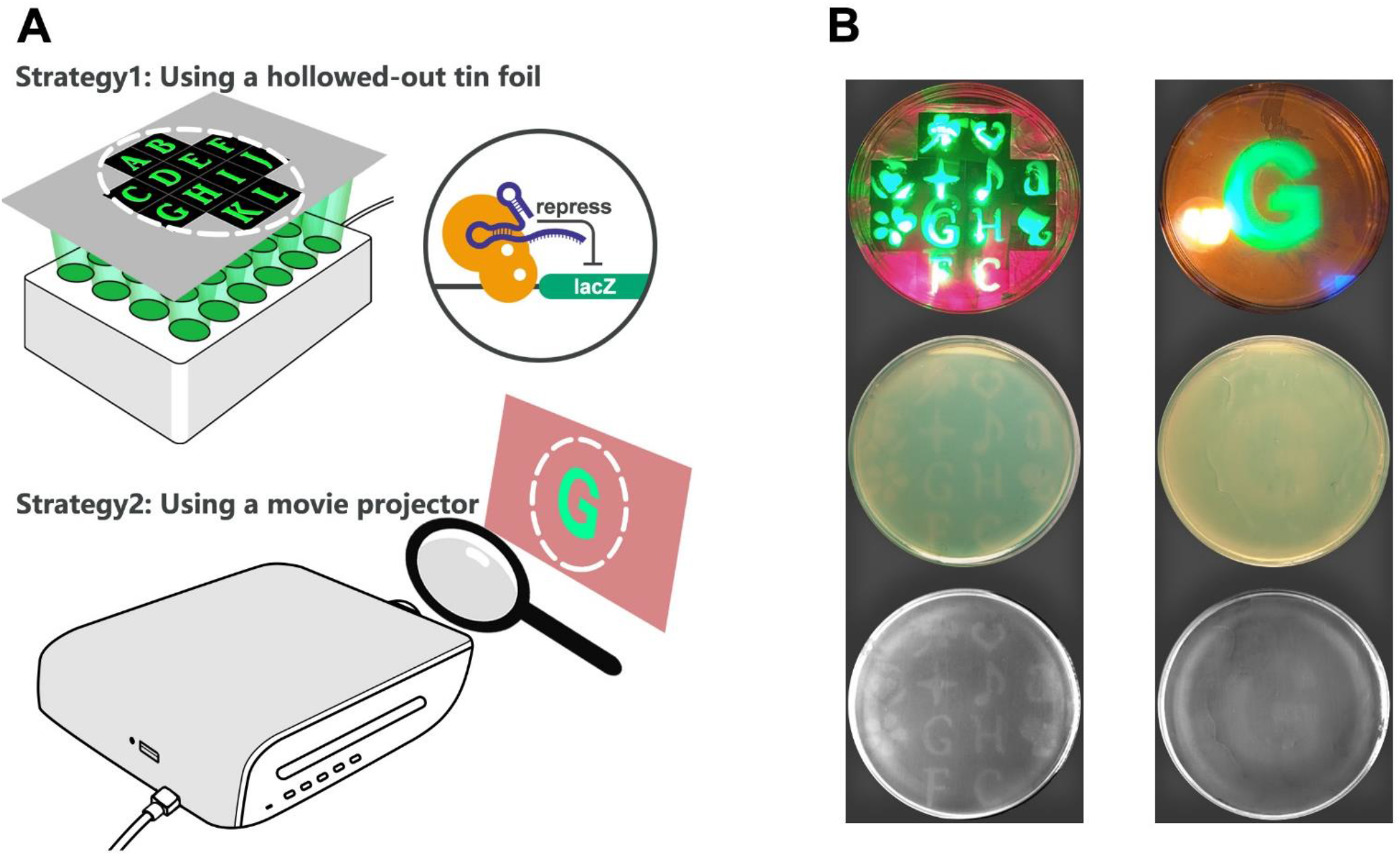
Two strategies were used to produce patterned *E. coli* plate. (A) An illustration of the working mechanism of optoCRISPRi-HD on *lacZ* and the devices used for light control. (B) Photographs that illustrated the light controlling effects on bacterial lawn of *E. coli* Nissle1917. Top: patterned green/red light illuminating *E. coli* culture plate that contained X-gal and IPTG. Middle: after 12 hours’ light control, blue-white pattern formed for both strategies, where green light illuminated regions remained uncolored. Bottom: photographs taken by a gel-imager for enhanced contrast.

### Light Controlled Filamentous Cell Formation by Repressing *ftsZ*

FtsZ is a tubulin homolog which is proven essential for cell division and the maintenance of cell morphology.^17–19^ In previous studies, the deficiency of FtsZ resulted in cell elongation^20,21^ while an over-expression caused smaller cells compared with normal.^22^ For targeting FtsZ, we co-transformed pvCq5 (carrying *ftsZ* sgRNA) and pvS108 into *E. coli* MGFROS strain. Cells were first cultivated in a red-light environment then transferred into a microfluidic chip, where they were constantly exposed to green light and recorded by live-cell imaging. As shown in **Figure 5**, the cells were normally shaped at time zero, but gradually became filamentous under green light (Supplementary video: optoCRISPRiHD-ftsZ.avi). After 4 hours’ exposure, *E. coli* were elongated significantly compared to the initial state, indicating the inability to form Z-ring caused by FtsZ deficiency under green light. This result suggested that FtsZ was normally expressed under red light then repressed under green light, which proved optoCRISPRi-HD is tight enough to regulate an essential gene.

**Figure 5.**
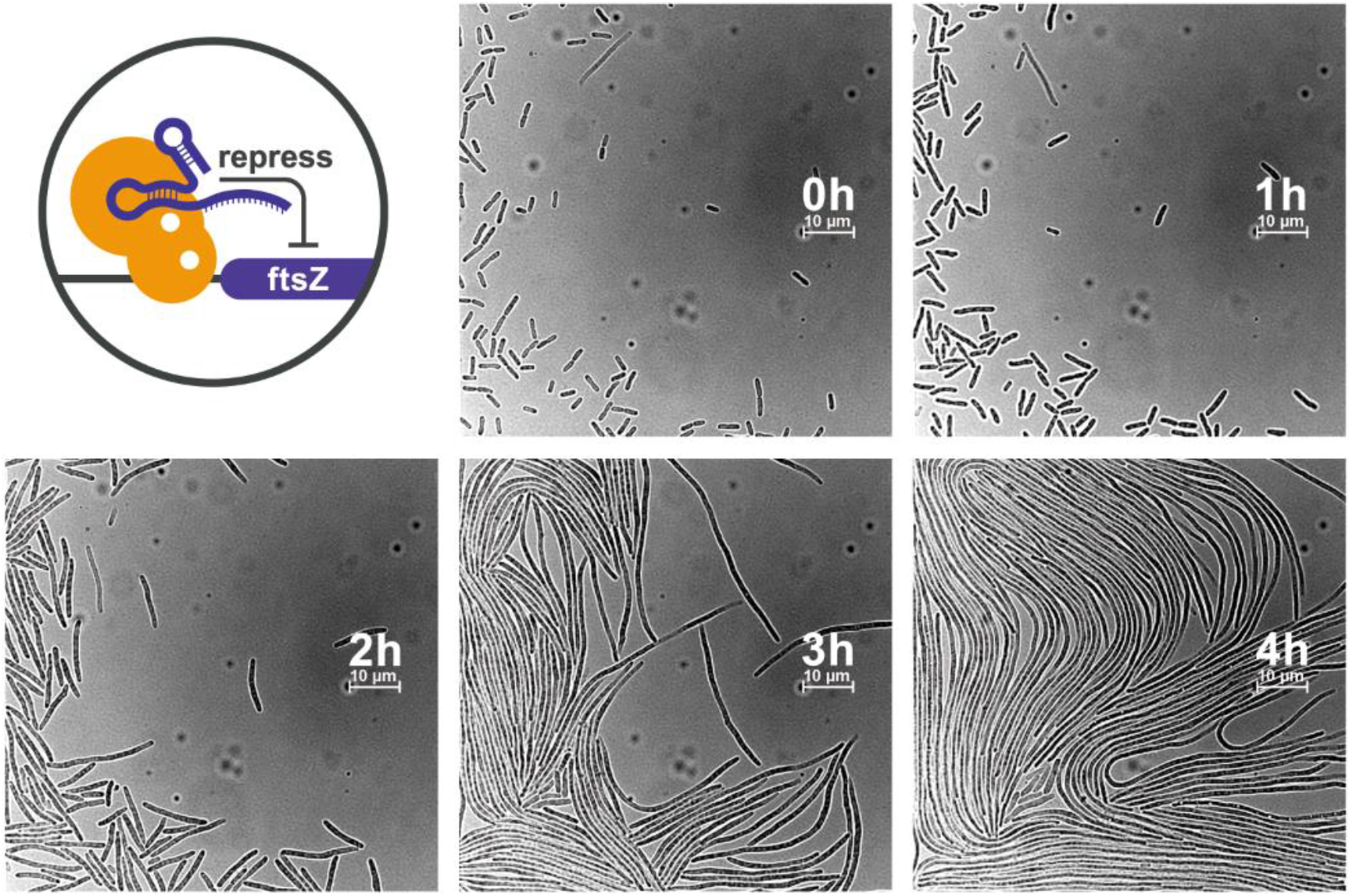
Microscopic images that record the continuous growth of the strain harboring pvS108 and pvCq5 under green light. The strain had grown in a red-light environment for 5 hours before time zero.

### Light Controlled Spherical Cell Formation by Repressing *mreB*

The actin homolog MreB is another essential protein that maintains cell shape. The depletion of MreB has been reported to cause cell enlargement and eventually cell death^23–25^ due to weakened cytoskeleton.^26,27^ Plasmid pvCq6 (carrying *mreB* sgRNA) was co-transformed with pvS108 into MGFROS strain to optically control MreB. Cells were cultivated in a red-light environment before being transferred into a microfluidic chip. As shown in **Figure 6**, after 4 hours’ exposure under green light, *E. coli* were significantly enlarged, the vast majority of which appeared to be ball-shaped instead of short rod-shaped (Supplementary video: optoCRISPRiHD-mreB.avi). This result suggested MreB was uninfluenced under red light and efficiently repressed under green light, another evidence to demonstrate the efficiency of optoCRISPRi-HD.

**Figure 6.**
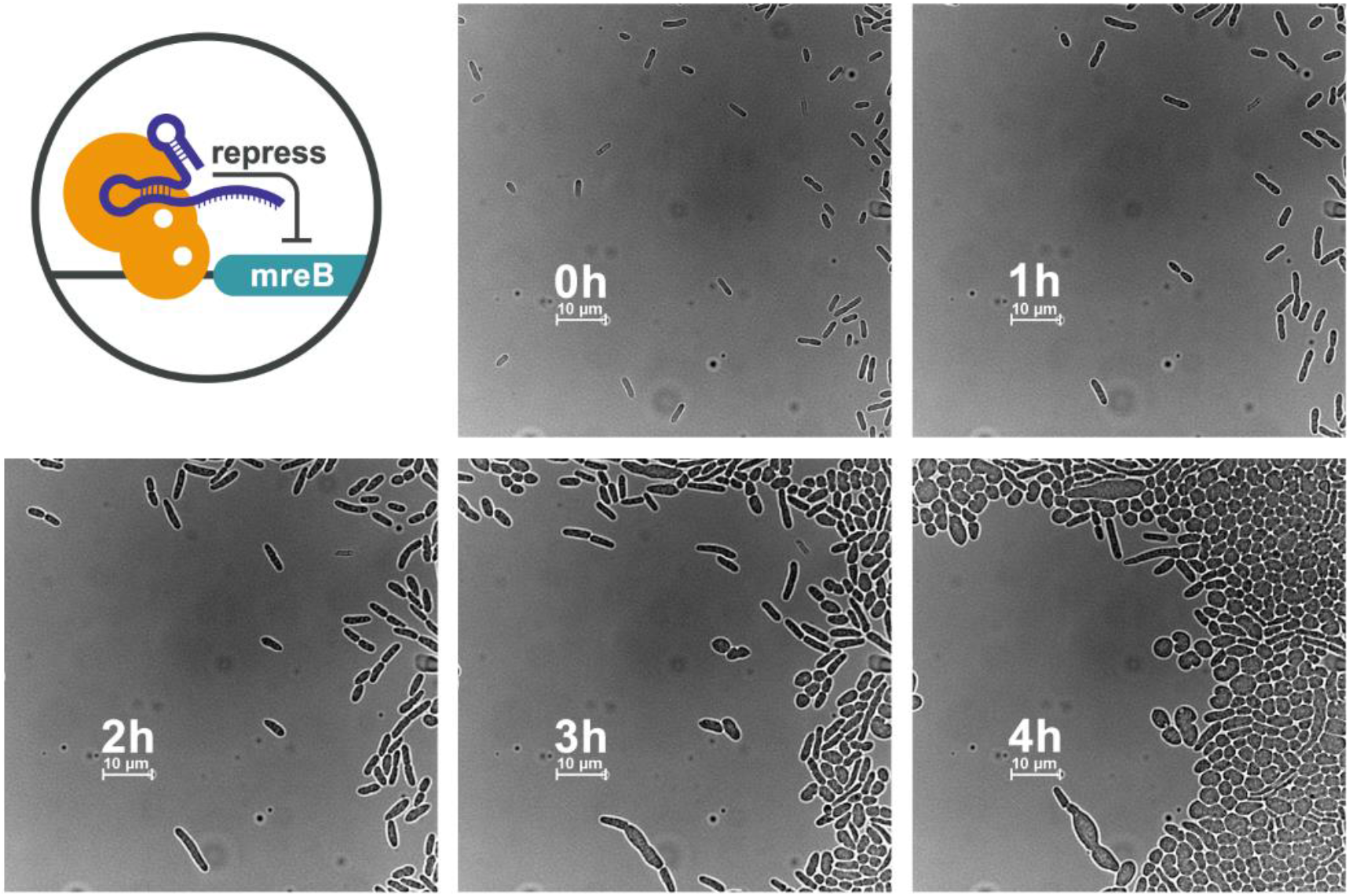
Microscopic images that record the continuous growth of the strain harboring pvS108 and pvCq6 under green light. The strain had grown in a red-light environment for 5 hours before time zero.

### Blocking the Initiation of DNA Replication with optoCRISPRi-HD

It has been reported that CRISPRi was able to block replication initiation.^28^ In *E. coli*, replication is initiated when DnaA proteins bind to DnaA boxes (specific sequences within *oriC*)^29^ and melt the adjacent AT-rich region to load DnaB. The occupation of dCas9 on DnaA boxes will disrupt DnaA binding and result in failed replication initiation.^28^ To test if optoCRISPRi-HD can achieve the same effect, we co-transformed plasmid pvCq3 (carrying *oriC* sgRNA) and pvS108 (**Figure S1**) into MGFROS strain. *E. coli* cells were first cultivated under red light then transferred into a microfluidic chip and illuminated by green light. After 4 hours’ green light exposure, arabinose was added into culture media for 1 hour to induce TetR-eGFP. The number of fluorescent spots per area indicated the density of *oriC*. It has been reported that the alternation of cell shape caused by FtsZ or MreB shortage will not influence replication initiation.^30^ As shown in **Figure 7**, by contrasting four green light treated strains, we found that the density of foci was much lower when optoCRISPRi-HD targeted *oriC* than that of *ftsZ, mreB* or no target, which suggested replication initiation has been disturbed in *oriC-sgRNA* strain.

**Figure 7.**
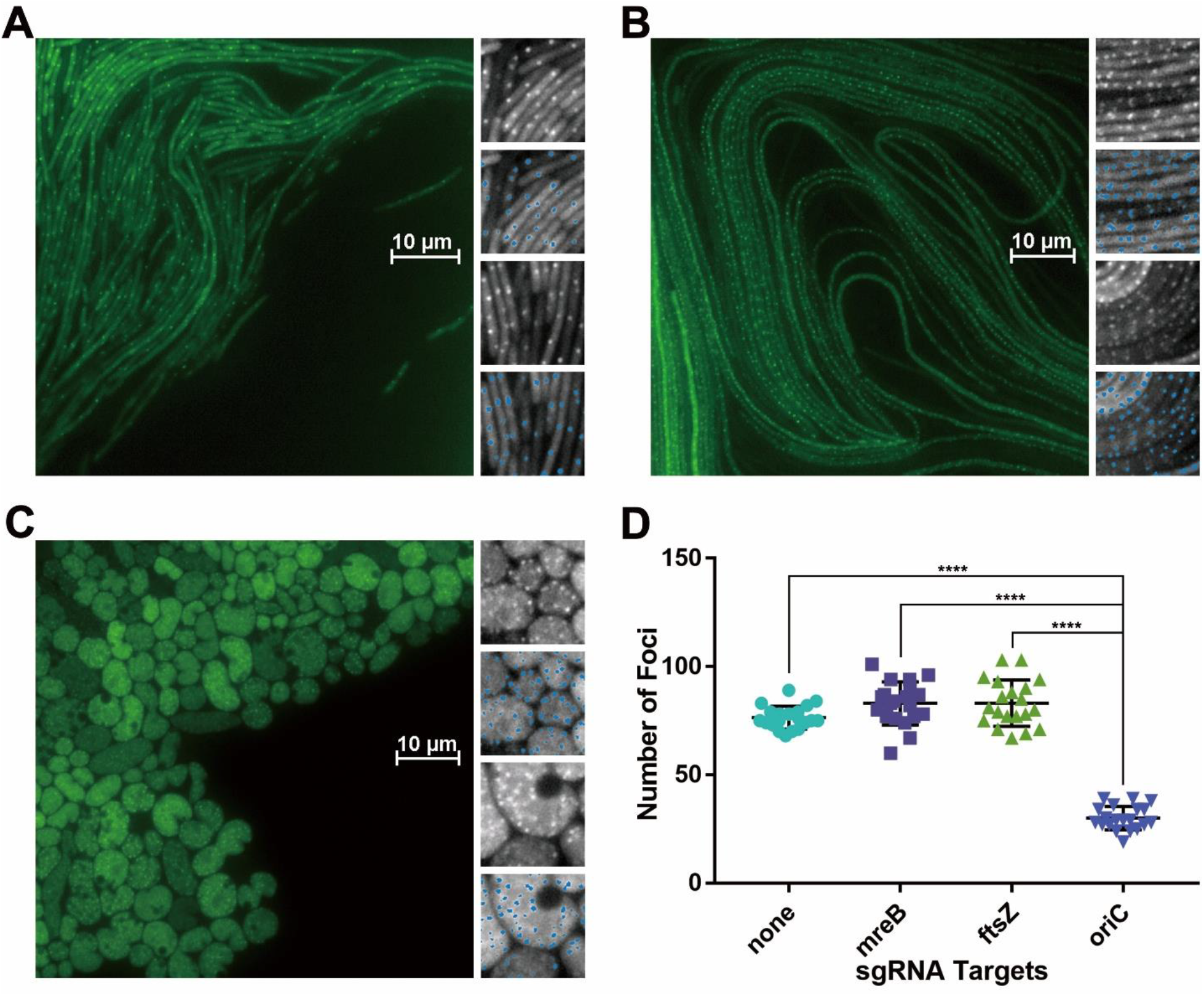
Statistics of *oriC* foci in green-light controlled strains. (A) Fluorescent images of *oriC* blocked strain. 100*100 pixel regions fulfilled by bacteria were chosen as sample unites to calculate the density of *oriC* foci. The difference of intercellular space between unites was neglected. Foci were distinguished by ilastik (Laplacian of Gaussian σ=1.0 and σ=1.6, blue spots). The number of foci was calculated by Fiji software. (B) Fluorescent images of FtsZ repressed strain. (C) Fluorescent images of MreB repressed strain. (D) Scatter plot of foci per area for different strains. 20 sample unites were counted for each group. The data of *oriC* blocked group showed significant difference compared with other groups (****: *p*-value = 0.0001, see **Table S3** for t-test details).

## DISCUSSION

Dynamic range is a major parameter to evaluate the performance of a molecular switch. A high dynamic range indicates strong effects when “switched on”and low leakage when “switched off”. Mathematic models^31,32^ and experiments^11^ suggested that CcaR level is crucial for enhanced dynamic range in CcaS-CcaR TCS. In this study, we tested a series of CcaR expression levels to find the best dynamic range for optoCRISPRi-HD. The result (**Figure 2**) showed that in this case, a higher dynamic range means lower leakage and limited total output of *P_cpcG2-172_* at the same time. For the control of essential genes, the priority of optoCRISPRi-HD is to minimize the leakage activity and ensure a normal cell growth before green light induction, which requires weak CcaR-RBSs like *J61108* and *J61110*. For other targets that require complete repression and have little effect on cell growth, stronger CcaR-RBS like *J61104* might be considered in future applications. Our experiments proved that *J23117-J61108-ccaR* is appropriate for balancing the needs of leakage control and repression effectiveness.

OptoCRISPRi-HD can be easily applied in bioprinting. Our movie projector experiment showed that despite the inaccuracy of green/red wavelengths, the digital light source was able to form a “G” pattern on culture plate. This result means a bioprinting can be achieved by our optoCRISPRi-HD without bandpass filters or patterned masks that were used in previous studies.^5,10,33^ We can assume that in future application scenarios like organ 3D printing, wavelengths and patterns are not fixed but able to change in real time, entirely controlled by computer programs.

Besides opto-CRISPRi, light controlled CRISPR^4,34^ and CRISPRa^5,35–37^ (CRISPR activation) systems have also been developed by researchers. Our study enriched the optogenetic toolbox and excavated the potential of optoCRISPRi-HD as a genome-wide gene regulation system with space-time precision and extensive targets. As a versatile tool with higher dynamic range than most, this system should facilitate further studies of gene networks, fermentation control or many other topics in synthetic biology.

## METHODS

### Reagents and Culture Media

Fluorescence quantification experiments were performed in EZ Rich Defined Medium (0.2% glucose or glycerol as carbon source). For all other experiments, LB Miller broth was used as standard medium for *E. coli* cultivation. Antibiotic concentrations were as follow: ampicillin sodium 20μg/ml, gentamycin sulfate 1.25μg/ml, kanamycin sulfate 10μg/ml, spectinomycin dihydrochloride 50μg/ml, chloramphenicol 25μg/ml.

In light controlled bioprinting experiments, 100μg/ml IPTG and 80μg/ml X-gal were added into LB plate. For fluorescence quantification, 0mM, 5mM, 10mM arabinose were added into RDM (glycerol as carbon source) respectively to induce genomic *tetR-eGFP*. 20mM arabinose was used to induce *oriC* foci for microscopic observation. All medium for microfluidic experiments contained 20mg/L Pluronic F127 surfactant (CAS 9003-11-6, Sigma-Aldrich).

### Construction of MGFROS Strain

We designed a 120 repeats’ *tetO* sequence to label *oriC*, in which every *tetO* was a 15bp sequence containing only the core region of TetR binding site^38^ and spaced by 10bp random sequences. The *tetO-GmR(gentamicin resistance gene)-tetO* sequence was made by DNA synthesis, along with the *asnA* and *viaA* overlaps. In order to obtain linearized fragment, the pUC57 plasmid containing the synthesized sequence was digested by BsaI and DraI. The FROS fragment was inserted into an *E. coli* MG1655 strain *(ΔenvZ ΔcheZ)* using λRed system. The gentamicin resistance is not removed from the genome of this strain. After the insertion of *tetO-GmR-tetO*, a *tetR-eGFP-loxP-Kan(kanamycin resistance gene)-loxP* sequence was inserted into genomic *ara* operon of the same strain. The kanamycin resistance was then removed from this strain. The final version of this strain is named MGFROS.

### Plasmid Assembly

ClonExpress (based on 15-20bp overlapping DNA ends, Vazyme C112-02 C113-01) was used as standard method of all plasmid assembly experiments. Plasmids and primers used are listed in **Table S1** and **Table S2**. For promoter and RBS changing experiments, the J23 or J61 plasmid series were obtained by reverse PCR. Plasmids pvCq1~7 (harboring different sgRNAs) were also obtained by reverse PCR.

### Light Devices

The construction and function of our 24-wall-plate light controlling device (device1, **Figure S2A**) was similar to the Light Plate Apparatus (LPA),^39^ except for separated green (525nm) or red (660nm) LEDs. Device2 (**Figure S2B**) was a 100 lumen LCD movie projector. Device1 and device2 were placed inside 37°C incubators during light inducing experiments.

For light control during microscopic observation, an annular LED device (device3) was placed between the light source of Nikon Eclipse Ti2-E and a homeothermic container of microfluidic chip (**Figure S2C**).

### Fluorescence Quantification

In promoter or RBS changing experiments, plasmids pvK0 series were co-transformed with pvC0 into MGFROS strain. Three biological repeats were isolated and cultivated overnight at 37°C, 200rpm as seed culture. The overnight seeds were added into RDM (glucose as carbon source) in a 24-wall plate at a ratio of 3‰. Two plates that contained identical samples were placed in green light device1 or red light device1 separately and cultivated at 37°C, 200rpm for 5 hours. The fluorescent result was measured using a microplate reader (Biotek Synergy H1).

In *tetR-eGFP* repressing experiment, plasmids pvS108 and pvCq1 (**Figure S1**) were co-transformed into MGFROS strain. Three biological repeats were isolated and cultivated overnight at 37°C, 200rpm as seed culture. The seeds were added into RDM (glycerol as carbon source, contained 0mM, 5mM,10mM arabinose respectively) in a 24-wall plate at a ratio of 3‰. Two plates that contained identical samples were placed in green light device1 or red light device1 separately and cultivated at 37°C, 200rpm for 8 hours. Fluorescent result was measured by the microplate reader described above.

### Microfluidics and Microscopy

For each experiment, we isolated a newly transformed colony to cultivate it in 15ml LB, red light environment (device1), 37°C, 200rpm for 5 hours. The culture was concentrated at 2500rpm, 12°C for 10 minutes to 0.5-0.6ml and injected into a microfluidic chip (**Figure S3**).

Microscopic observation was performed using Nikon Eclipse Ti2-E (100x oil immersion objective). The microfluidic chip was placed in a homeothermic container with a transparent top and illuminated by device3 (**Figure S2C**) for 4 hours, during which the image of bacteria was taken every 30 seconds through PFS (perfect focus system) and adjusted by automatic LUT. After 4 hours’ green light exposure, the original LB media was taken place by LB that contained 20mM arabinose to induce *oriC* foci. The fluorescent images were taken 1 hour after arabinose induction.

## Supporting information

optoCRISPRiHD-ftsZ-video

optoCRISPRiHD-mreB-video

## ASSOCIATED CONTENT

Supplementary materials: supplementary figures and tables.

Supplementary videos

## ACKNOWLEDGEMENTS

This work was supported by the National Natural Science Foundation of China (Grant 31971184). The funders had no role in study design, data collection and interpretation, or the decision to submit the work for publication. Help from Chun-Xiong Luo (School of Physics, Peking University) and Hui-Min Qi (former member of our lab) is highly appreciated.

## REFERENCES

(1) Qi, L. S., Larson, M. H., Gilbert, L. A., Doudna, J. A., Weissman, J. S., Arkin, A. P., & Lim, W. A. (2013). Repurposing CRISPR as an RNA-guided platform for sequence-specific control of gene expression. Cell, 152(5), 1173–1183.

(2) Cho, S., Choe, D., Lee, E., Kim, S. C., Palsson, B., & Cho, B. K. (2018). High-Level dCas9 Expression Induces Abnormal Cell Morphology in Escherichia coli. ACS synthetic biology, 7(4), 1085–1094.

(3) Wu, H., Wang, Y., Wang, Y., Cao, X., Wu, Y., Meng, Z., Su, Q., Wang, Z., Yang, S., Xu, W., Liu, S., Cheng, P., Wu, J., Khan, M. R., He, L., & Ma, G. (2014). Quantitatively relating gene expression to light intensity via the serial connection of blue light sensor and CRISPRi. ACS synthetic biology, 3(12), 979–982.

(4) Nihongaki, Y., Kawano, F., Nakajima, T., & Sato, M. (2015). Photoactivatable CRISPR-Cas9 for optogenetic genome editing. Nature biotechnology, 33(7), 755–760.

(5) Fernandez-Rodriguez, J., Moser, F., Song, M., & Voigt, C. A. (2017). Engineering RGB color vision into Escherichia coli. Nature chemical biology, 13(7), 706–708.

(6) Wu, P., Chen, Y., Liu, M., Xiao, G., & Yuan, J. (2021). Engineering an Optogenetic CRISPRi Platform for Improved Chemical Production. ACS synthetic biology, 10(1), 125–131.

(7) Liang, J. Y., Yuann, J. M., Cheng, C. W., Jian, H. L., Lin, C. C., & Chen, L. Y. (2013). Blue light induced free radicals from riboflavin on E. coli DNA damage. Journal of photochemistry and photobiology. B, Biology, 119, 60–64.

(8) Liang, J. Y., Cheng, C. W., Yu, C. H., & Chen, L. Y. (2015). Investigations of blue light-induced reactive oxygen species from flavin mononucleotide on inactivation of E. coli. Journal of photochemistry and photobiology. B, Biology, 143, 82–88.

(9) Dong, K., Goyarts, E. C., Pelle, E., Trivero, J., & Pernodet, N. (2019). Blue light disrupts the circadian rhythm and create damage in skin cells. International journal of cosmetic science, 41(6), 558–562.

(10) Tabor, J. J., Levskaya, A., & Voigt, C. A. (2011). Multichromatic control of gene expression in Escherichia coli. Journal of molecular biology, 405(2), 315–324.

(11) Schmidl, S. R., Sheth, R. U., Wu, A., & Tabor, J. J. (2014). Refactoring and optimization of light-switchable Escherichia coli two-component systems. ACS synthetic biology, 3(11), 820–831.

(12) Gambetta, G. A., & Lagarias, J. C. (2001). Genetic engineering of phytochrome biosynthesis in bacteria. Proceedings of the National Academy of Sciences of the United States of America, 98(19), 10566–10571.

(13) Nakajima, M., Ferri, S., Rögner, M., & Sode, K. (2016). Construction of a Miniaturized Chromatic Acclimation Sensor from Cyanobacteria with Reversed Response to a Light Signal. Scientific reports, 6, 37595.

(14) Ong, N. T., & Tabor, J. J. (2018). A Miniaturized Escherichia coli Green Light Sensor with High Dynamic Range. Chembiochem: a European journal of chemical biology, 19(12), 1255–1258.

(15) Hirose, Y., Shimada, T., Narikawa, R., Katayama, M., & Ikeuchi, M. (2008). Cyanobacteriochrome CcaS is the green light receptor that induces the expression of phycobilisome linker protein. Proceedings of the National Academy of Sciences of the United States of America, 105(28), 9528–9533.

(16) Lau, I. F., Filipe, S. R., Søballe, B., Økstad, O. A., Barre, F. X., & Sherratt, D. J. (2003). Spatial and temporal organization of replicating Escherichia coli chromosomes. Molecular microbiology, 49(3), 731–743.

(17) Margolin W. (2005). FtsZ and the division of prokaryotic cells and organelles. Nature reviews. Molecular cell biology, 6(11), 862–871.

(18) Hill, N. S., Kadoya, R., Chattoraj, D. K., & Levin, P. A. (2012). Cell size and the initiation of DNA replication in bacteria. PLoS genetics, 8(3), e1002549.

(19) de Boer, P. A., Crossley, R. E., & Rothfield, L. I. (1992). Roles of MinC and MinD in the site-specific septation block mediated by the MinCDE system of Escherichia coli. Journal of bacteriology, 174(1), 63–70.

(20) Bi, E., & Lutkenhaus, J. (1993). Cell division inhibitors SulA and MinCD prevent formation of the FtsZ ring. Journal of bacteriology, 175(4), 1118–1125.

(21) Chien, A. C., Hill, N. S., & Levin, P. A. (2012). Cell size control in bacteria. Current biology: CB, 22(9), R340–R349.

(22) Ward, J. E., Jr, & Lutkenhaus, J. (1985). Overproduction of FtsZ induces minicell formation in E. coli. Cell, 42(3), 941–949.

(23) Alyahya, S. A., Alexander, R., Costa, T., Henriques, A. O., Emonet, T., & Jacobs-Wagner, C. (2009). RodZ, a component of the bacterial core morphogenic apparatus. Proceedings of the National Academy of Sciences of the United States of America, 106(4), 1239–1244.

(24) van den Ent, F., Johnson, C. M., Persons, L., de Boer, P., & Löwe, J. (2010). Bacterial actin MreB assembles in complex with cell shape protein RodZ. The EMBO journal, 29(6), 1081–1090.

(25) van den Ent, F., Amos, L. A., & Löwe, J. (2001). Prokaryotic origin of the actin cytoskeleton. Nature, 413(6851), 39–44.

(26) Jiang, X. R., & Chen, G. Q. (2016). Morphology engineering of bacteria for bio-production. Biotechnology advances, 34(4), 435–440.

(27) Bendezú, F. O., Hale, C. A., Bernhardt, T. G., & de Boer, P. A. (2009). RodZ (YfgA) is required for proper assembly of the MreB actin cytoskeleton and cell shape in E. coli. The EMBO journal, 28(3), 193–204.

(28) Wiktor, J., Lesterlin, C., Sherratt, D. J., & Dekker, C. (2016). CRISPR-mediated control of the bacterial initiation of replication. Nucleic acids research, 44(8), 3801–3810.

(29) Yung, B. Y., & Kornberg, A. (1989). The dnaA initiator protein binds separate domains in the replication origin of Escherichia coli. The Journal of biological chemistry, 264(11), 6146–6150.

(30) Zheng, H., Ho, P. Y., Jiang, M., Tang, B., Liu, W., Li, D., Yu, X., Kleckner, N. E., Amir, A., & Liu, C. (2016). Interrogating the Escherichia coli cell cycle by cell dimension perturbations. Proceedings of the National Academy of Sciences of the United States of America, 113(52), 15000–15005.

(31) Nishida, A., Sekine, R., Kiga, D., & Yamamura, M. (2016). High-frequency noise attenuation of a two-component system responding to short-pulse input. In Proceedings of the 2016 7th International Conference on Computational Systems-Biology and Bioinformatics, CSBio 2016 (pp. 28–35).

(32) Olson, E. J., Tzouanas, C. N., & Tabor, J. J. (2017). A photoconversion model for full spectral programming and multiplexing of optogenetic systems. Molecular systems biology, 13(4), 926.

(33) Tabor, J. J., Salis, H. M., Simpson, Z. B., Chevalier, A. A., Levskaya, A., Marcotte, E. M., Voigt, C. A., & Ellington, A. D. (2009). A synthetic genetic edge detection program. Cell, 137(7), 1272–1281.

(34) Yu, Y., Wu, X., Guan, N., Shao, J., Li, H., Chen, Y., Ping, Y., Li, D., & Ye, H. (2020). Engineering a far-red light-activated split-Cas9 system for remote-controlled genome editing of internal organs and tumors. Science advances, 6(28), eabb1777.

(35) Nihongaki, Y., Yamamoto, S., Kawano, F., Suzuki, H., & Sato, M. (2015). CRISPR-Cas9-based photoactivatable transcription system. Chemistry & biology, 22(2), 169–174.

(36) Polstein, L. R., & Gersbach, C. A. (2015). A light-inducible CRISPR-Cas9 system for control of endogenous gene activation. Nature chemical biology, 11(3), 198–200.

(37) Nihongaki, Y., Furuhata, Y., Otabe, T., Hasegawa, S., Yoshimoto, K., & Sato, M. (2017). CRISPR-Cas9-based photoactivatable transcription systems to induce neuronal differentiation. Nature methods, 14(10), 963–966.

(38) Orth, P., Schnappinger, D., Hillen, W., Saenger, W., & Hinrichs, W. (2000). Structural basis of gene regulation by the tetracycline inducible Tet repressor-operator system. Nature structural biology, 7(3), 215–219.

(39) Gerhardt, K. P., Olson, E. J., Castillo-Hair, S. M., Hartsough, L. A., Landry, B. P., Ekness, F., Yokoo, R., Gomez, E. J., Ramakrishnan, P., Suh, J., Savage, D. F., & Tabor, J. J. (2016). An open-hardware platform for optogenetics and photobiology. Scientific reports, 6, 35363.

