## Supplementary figures and tables for "OptoCRISPRi-HD: engineering a green-light activated CRISPRi system with high dynamic range"

### Supplementary Materials

#### Supplementary Figures

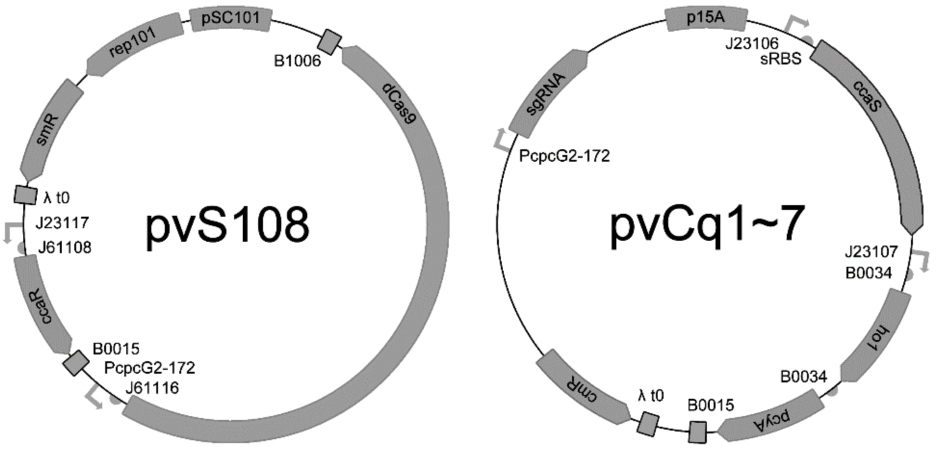


**Figure S1.** Plasmids pvS108 and pvCq1~7.

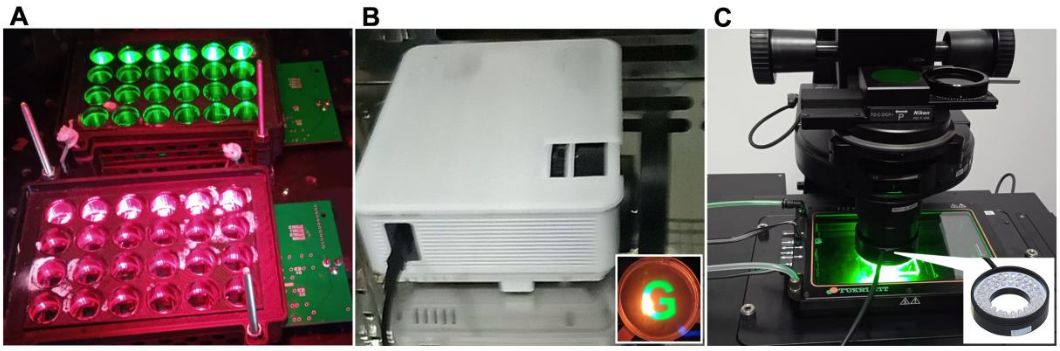


**Figure S2.** Light devices used in this study. (A) Device1 was built based on Light Plate

Apparatus (LPA)[1]. The core of device1 was a printed circuit board (PCB). (B) Device2 was a 100 lumen LCD movie projector. A 10-fold magnifier was used to shorten the imaging distance. (C) Device3 was an annular LED machine-vision light source. The LEDs were arranged on a slightly beveled surface to focus the light.

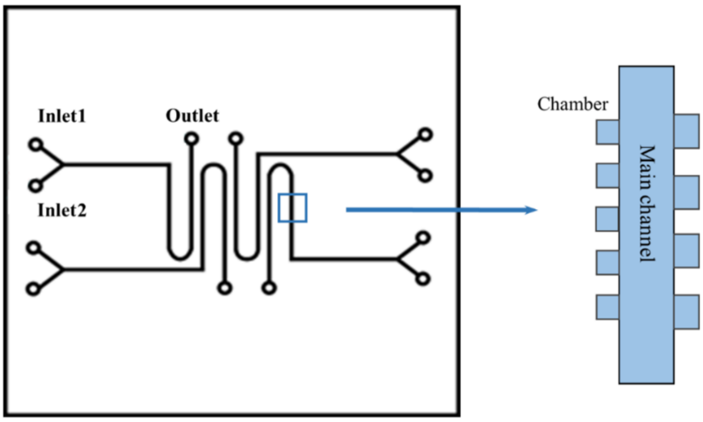


**Figure S3.** Layout of microfluidic chips used in this study. The height of each chamber is 1μm.

### Supplementary Tables

**Table S1.** Plasmids constructed in this study

| Name | Replicon/Resistance | Contents |
|---|---|---|
| pvS108 | pSC101/SmR | *ccaR*; *dCas9* |
| pvCq1 | p15A/CmR | *ccaS*; *ho1*; *pcyA*; *sgRNA* (targets *tetR-eGFP*) |
| pvCq2 | p15A/CmR | *ccaS*; *ho1*; *pcyA*; *sgRNA* (targets *lacZ* in MGFROS) |
| pvCq3 | p15A/CmR | *ccaS*; *ho1*; *pcyA*; *sgRNA* (targets *oriC*) |
| pvCq4 | p15A/CmR | *ccaS*; *ho1*; *pcyA*; *sgRNA* (targets *dnaA*) |
| pvCq5 | p15A/CmR | *ccaS*; *ho1*; *pcyA*; *sgRNA* (targets *ftsZ*) |
| pvCq6 | p15A/CmR | *ccaS*; *ho1*; *pcyA*; *sgRNA* (targets *mreB*) |
| pvCq7 | p15A/CmR | *ccaS*; *ho1*; *pcyA*; *sgRNA* (targets *lacZ* in Nissle1917) |
| pvK0.0.00 | pSC101/KanR | *ccaR (J23100-J61100)* |
| pvK0.9.00 | pSC101/KanR | *ccaR (J23109-J61100)*; *eGFP* |
| pvK0.4.00 | pSC101/KanR | *ccaR (J23114-J61100)*; *eGFP* |
| pvK0.5.00 | pSC101/KanR | *ccaR (J23115-J61100)*; *eGFP* |
| pvK0.7.00 | pSC101/KanR | *ccaR (J23117-J61100)*; *eGFP* |
| pvK0.7.04 | pSC101/KanR | *ccaR (J23117-J61104)*; *eGFP* |
| pvK0.7.08 | pSC101/KanR | *ccaR (J23117-J61108)*; *eGFP* |
| pvK0.7.10 | pSC101/KanR | *ccaR (J23117-J61110)*; *eGFP* |
| pvK0.7.12 | pSC101/KanR | *ccaR (J23117-J61112)*; *eGFP* |
| pvK0.7.14 | pSC101/KanR | *ccaR (J23117-J61114)*; *eGFP* |
| pvK0.7.16 | pSC101/KanR | *ccaR (J23117-J61116)*; *eGFP* |
| pvC0 | p15A/CmR | *ccaS*; *ho1*; *pcyA* |

**Notes: (1) About plasmids pvCq1~7.** Compared to CcaS-CcaR v2.0[2], we deleted the B0015 terminator between *ccaS* and *ho1* to increase the stability of pvCq1~7 and reduce the size of the plasmids. The original *J23106* promoter for *ho1* and *pcyA* was replaced by *J23107*. **(2) About pvCq4 that targets *dnaA*.** We designed pvCq4 that targets *dnaA* in order to observe failed initiation of chromosome DNA replication under green light. This effect was unstable among tests, possibly because DnaA is also necessary for the initiation of plasmid DNA replication, thus the expression of dCas9 and sgRNA would cause trouble for their own plasmids to maintain in *E. coli* cells.

**Table S2.** Primers used to change sgRNA

| Sequence | Use |
|---|---|
| Q1-F: aatagcttctgggcgagtttacgttttagagctagaaatagcaagttaaaataaggc | Acquiring |
| Q1-R: aaactcgcccagaagctattaaaaatgcgatcctaacaaagtaaaattgaagaaaag | pvCq1 |
| Q2-F: aaggccagtgaatccgtaatcagttttagagctagaaatagcaagttaaaataaggc | Acquiring |
| Q2-R: attacggattcactggccttaaaaatgcgatcctaacaaagtaaaattgaagaaaag | pvCq2 |
| Q3-F: aactactgtggataactctgtcgttttagagctagaaatagcaagttaaaataaggc | Acquiring |
| Q3-R: cagagttatccacagtagttaaaaatgcgatcctaacaaagtaaaattgaagaaaag | pvCq3 |
| Q4-F: aatggtaactcatcctgcaatcgttttagagctagaaatagcaagttaaaataaggc | Acquiring |
| Q4-R: ttgcaggatgagttaccattaaaaatgcgatcctaacaaagtaaaattgaagaaaag | pvCq4 |
| Q5-F: aagactttaatcaccgcgtcatgttttagagctagaaatagcaagttaaaataaggc | Acquiring |
| Q5-R: gacgcggtgattaaagtcttaaaaatgcgatcctaacaaagtaaaattgaagaaaag | pvCq5 |
| Q6-F: aaattcgcagtacccaggtcaagttttagagctagaaatagcaagttaaaataaggc | Acquiring |
| Q6-R: gacctgggtactgcgaatttaaaaatgcgatcctaacaaagtaaaattgaagaaaag | pvCq6 |
| Q7-F: aagtcacgacgttgtaatacgagttttagagctagaaatagcaagttaaaataaggc | Acquiring |
| Q7-R: gtattacaacgtcgtgacttaaaaatgcgatcctaacaaagtaaaattgaagaaaag | pvCq7 |

**Table S3.** T-test analysis of *oriC* targeting group compared to other groups

| Dunnett's multiple comparisons test | Mean Diff. | 95.00% CI of diff. | Significant? | Summary | Adjusted P-Value | D-? | | |
|---|---|---|---|---|---|---|---|---|
| oriC vs. none | -46.4 | [-51.06, -41.74] | Yes | **** | 0.0001 | A | none | |
| oriC vs. mreB | -53 | [-59.34, -46.66] | Yes | **** | 0.0001 | B | mreB | |
| oriC vs. ftsZ | -53.1 | [-60.13, -46.07] | Yes | **** | 0.0001 | C | ftsZ | |
| Test details | Mean 1 | Mean 2 | Mean Diff. | SE of diff. | n1 | n2 | q | DF |
| oriC vs. none | 30 | 76.4 | -46.4 | 1.826 | 20 | 20 | 25.41 | 19 |
| oriC vs. mreB | 30 | 83 | -53 | 2.486 | 20 | 20 | 21.32 | 19 |
| oriC vs. ftsZ | 30 | 83.1 | -53.1 | 2.754 | 20 | 20 | 19.28 | 19 |

### References


(1) Gerhardt, K. P., Olson, E. J., Castillo-Hair, S. M., Hartsough, L. A., Landry, B. P., Ekness, F., Yokoo, R., Gomez, E. J., Ramakrishnan, P., Suh, J., Savage, D. F., & Tabor, J. J. (2016). An open-hardware platform for optogenetics and photobiology. *Scientific reports*, *6*, 35363.

(2) Schmidl, S. R., Sheth, R. U., Wu, A., & Tabor, J. J. (2014). Refactoring and optimization of light-switchable Escherichia coli two-component systems. *ACS synthetic biology*, *3*(11), 820–831.
